# Frequent expansion of *Plasmodium vivax* Duffy Binding Protein in Ethiopia and its epidemiological significance

**DOI:** 10.1101/543959

**Authors:** Eugenia Lo, Jessica B. Hostetler, Delenasaw Yewhalaw, Richard D. Pearson, Muzamil M. A. Hamid, Karthigayan Gunalan, Daniel Kepple, Anthony Ford, Daniel A. Janies, Julian C. Rayner, Louis H. Miller, Guiyun Yan

**Author notes:** Correspondence authors: Eugenia Lo < >; Guiyun Yan< >; Louis H. Miller < >.

## Abstract

*Plasmodium vivax* invasion of human erythrocytes depends on the Duffy Binding Protein (*PvDBP*) which interacts with the Duffy antigen. *PvDBP* copy number varies between *P. vivax* isolates, but the prevalence of *PvDBP* multiplications in Sub-Saharan Africa and its impact are unknown. We determined the prevalence and type of *PvDBP* duplications, as well as *PvDBP* copy number variation among 178 Ethiopian *P. vivax* isolates using a PCR-based diagnostic method, a novel quantitative real-time PCR assay and whole genome sequencing. For the 145 symptomatic samples, *PvDBP* duplications were detected in 95 isolates, of which 81 had the Cambodian and 14 Malagasy-type *PvDBP* duplications. *PvDBP* varied from 1 to >4 copies. Isolates with multiple *PvDBP* copies were found to be higher in symptomatic than asymptomatic infections. For the 33 asymptomatic samples, *PvDBP* was detected with two copies in two of the isolates, and both were the Cambodian-type *PvDBP* duplication. *PvDBP* copy number in Duffy-negative heterozygotes was not significantly different from that in Duffy-positives, providing no support for the hypothesis that increased copy number is a specific association with Duffy-negativity, although the number of Duffy-negatives was small and further sampling is required to test this association thoroughly.

**Author summary:** *Plasmodium vivax* invasion of human erythrocytes relies on interaction between the Duffy antigen and *P. vivax* Duffy Binding Protein (*PvDBP*). Whole genome sequences from *P. vivax* field isolates in Madagascar identified a duplication of the *PvDBP* gene and *PvDBP* duplication has also been detected in non-African *P. vivax*-endemic countries.

Two types of *PvDBP* duplications have been reported, termed Cambodian and Malagasy-type duplications. Our study used a combination of PCR-based diagnostic method, a novel quantitative real-time PCR assay, and whole genome sequencing to determine the prevalence and type of *PvDBP* duplications, as well as *PvDBP* copy number on a broad number of *P. vivax* samples in Ethiopia. We found that over 65% of *P. vivax* isolated from the symptomatic infections were detected with *PvDBP* duplications and *PvDBP* varied from 1 to >4 copies. The majority of *PvDBP* duplications belongs to the Cambodian-type while the Malagasy-type duplications was also detected. For the asymptomatic infections, despite a small sample size, the majority of *P. vivax* were detected with a single-copy based on both PCR and qPCR assays. There was no significant difference in *PvDBP* copy number between Duffy-null heterozygote and Duffy-positive homozygote/heterozygote. Further investigation is needed with expanded Duffy-null homozygotes to examine the functional significance of *PvDBP* expansion.

## Introduction

*Plasmodium vivax* and *P. falciparum* are the two major parasite species that cause clinical malaria worldwide. While *P. falciparum* causes most malaria mortality, *P. vivax* can also cause severe disease and, more rarely death [1,2]. *P. vivax* is prominent in India, Southeast Asia, and South America. By contrast, vivax malaria is rare in sub-Saharan Africa, with significant rates of infection only found in a few countries [3]. The rarity of this infection is tightly associated with high prevalence of Duffy negativity in West and Central Africa [4]. This association represents a classic example of innate resistance to a pathogen resulting from the absence of a receptor for the pathogen [5]. A point mutation in the GATA-1 transcription factor binding site of the Duffy antigen/receptor for chemokines (*DARC*) gene promoter eliminates Duffy antigen (Fy) expression on the RBC surface required for *P. vivax* invasion [6]. This phenomenon is thought to account for the lack of *P. vivax* malaria in sub-Saharan Africa.

In Ethiopia, *P. vivax* and *P. falciparum* account for approximately 40% and 60% of the malaria cases, respectively [7,8]. The proportion of Duffy negatives in Ethiopia is about 35% [9,10], considerably lower than that documented in West and Central Africa (>97%) [4]. About 20% of hospital patients presented with malaria symptoms are Duffy-negatives [10–11]. While Duffy negativity was previously thought to be associated with complete resistance to *P. vivax*, *P. vivax* infection in Duffy-negative individuals was confirmed unequivocally in Madagascar [12], and has also been observed in multiple other parts of Africa as well as other parts of the world [13–18].

*Plasmodium vivax* invasion of human erythrocytes relies on interaction between the Duffy antigen and *P. vivax* Duffy Binding Protein (*PvDBP*) [19–20], though other reticulocyte ligand-receptor have also been recently identified [21]. Our recent study has shown that mutations in *PvDBP* do not explain the ability of *P. vivax* parasites to infect Duffy-negative individuals [22], but there could be other molecular mechanisms at play. Whole genome sequences from *P. vivax* field isolates in Madagascar identified a duplication of the *PvDBP* gene [23] and *PvDBP* duplication has also been detected in non-African *P. vivax*-endemic countries [24]. The high prevalence of *PvDBP* duplications raises the possibility that it is linked to the ability of *P. vivax* to infect Duffy-negative individuals. Two types of *PvDBP* duplications have been reported, termed Cambodian and Malagasy-type duplications based on the samples in which they were first described [23–24]. Both create a complete duplication of the *PvDBP* gene, but they differ in the sequence and length of the 3’-boundary of the linker region between the tandem copies of *PvDBP* [24]. In this study, we used whole genome sequencing and PCR-based diagnostic methods to determine the prevalence of both the Madagascar- and Cambodia-type *PvDBP* duplications in southwestern Ethiopia where a high incidence of *P. vivax* infections and a modest proportion of Duffy-null individuals co-occur. We compared *PvDBP* copy number between Duffy-null heterozygotes (*FyA*/*B^ES^* or*FyB*/*B^ES^*) and homozygotes (*FyB^ES^*/*B^ES^*), between symptomatic and asymptomatic infections, and explored the correlation of gene copy number with parasitemia, age, gender, ethnicity, and malaria symptoms in order to shed light on the epidemiological significance of *PvDBP* duplications.

## Materials and Methods

### Ethics statement

Scientific and ethical clearance was given by the institutional scientific and ethical review boards of the Jimma University, Ethiopia, and University of North Carolina at Charlotte, USA. Written informed consent/assent for study participation was obtained from all consenting heads of households, parents/guardians (for minors under age of 18), and each individual who was willing to participate in the study. All experimental protocols were reviewed and approved by the institutional review board (IRB) of the University of North Carolina at Charlotte, USA and performed in accordance with the IRB guidelines and regulations.

### Sample collection

A total of 178 *P. vivax* samples were included in this study. They were isolated from 145 symptomatic patients and 33 asymptomatic volunteers in Jimma, Ethiopia that had previously been confirmed by microscopy and PCR to be infected with *P. vivax* [9,10]. Symptomatic samples were obtained from patients who visited the health centers/hospitals in the Jimma town and presented with malaria signs/symptoms including axillary temperature ≥37°C, vomiting, diarrhea, nausea, abdominal pain, chills, fatigue, muscle pain, headache, malaise or loss of appetite (Supplementary Table 1). The asymptomatic samples were obtained from the community in the same area and had no fever or other malaria symptoms at the time of collection. Thick and thin blood smears were prepared for each subject to screen for *P. vivax* by microscopy. Parasites were counted against 200 leukocytes and a slide was considered negative when no parasites were observed after counting over 100 microscopic fields. All slides were read in duplicate by two microscopists at the time of sample collection. The density of parasitemia was expressed as the number of asexual *P. vivax* per microliter of blood, assuming a leukocyte count of 8000 per microliter. Three to four blood spots, each equivalent to ~50μl of blood, were blotted on Whatman 3MM filter paper from each participating individual. Parasite DNA was extracted from dried blood spots by the Saponin/Chelexmethod [25] and genomic DNA was eluted in a total volume of 200 μl TE buffer. Eluted DNA was confirmed with *P. vivax* by nested and quantitative PCR using the 18S rRNA *P. vivax*-specific primers [10], prior to PCR diagnosis of *PvDBP* duplications and quantification of copy number.

For a subset of 20 symptomatic samples that were confirmed with *P. vivax*, 6ml whole blood was collected from the patients. We used the Lymphoprep/Plasmodipur-based protocol to deplete white cells and enrich red cells before DNA extraction [26]. DNA was extracted from the enriched RBCs using the ZymoBead Genomic DNA kit (Zymo Research) following the manufacturer’s procedures. Whole genome sequences were obtained for these samples on a HiSeq Illumina Sequencing Platform at the Wellcome Trust Sanger Institute (ENA accession number of each sample in Table 1). The generated sequence reads were mapped individually to the *P*. *vivax* PVP01 reference genome [27] using bwa version 0.5.9-r16 with default parameters [28]. The resulting assembly bam files were reviewed in the region containing *PvDBP* (chromosome 6: 976329–980090) using the Artemis genome viewer [29]. Using a “non-proper pair” read filter in Artemis, mate pairs that were oriented tail-to-tail indicated *PvDBP* duplications [24].

**Table 1.** Information of whole genome sequences of 20 *Plasmodium vivax* isolates from Ethiopia. The European Nucleotide Archive (ENA) accession number for all files (ERP012186).

| Blood sample | ENA accession number | Whole genome sequence |  |  |  |  |  |  |  | PCR diagnosis |  | qPCR |
| --- | --- | --- | --- | --- | --- | --- | --- | --- | --- | --- | --- | --- |
|  |  | All reads | *P. vivax* reads (%) | Average genome coverage | *PvDBP* coverage | *PvDBP* copy | Mate/Cvg support § (Yes/No) | Duplication type [23,24] (Cambodian/Malagasy) | Genome covered 15X (%) | Malagasy duplication BF+AR (Yes/No)* | Cambodia duplication BF+AR2 (Yes/No)* | GCN |
| BBH(1)-125 | ERS849384 | 16104099 | 14765167 (92%) | 83.11 | 352.9 | 4.25 | Yes | Cambodian | 99.6 | No | Yes | 4.30 |
| BBH(1)-132 | ERS2593850 | 295443478 | 269499723 (91%) | 582.99 | 548.8 | 0.94 | No | Single copy | 98.5 | No | No | 1.14 |
| BBH(1)-137 | ERS2593862 | 15428892 | 12612040 (82%) | 41.09 | 99.14 | 2.41 | Yes | Cambodian | 96.1 | No | Yes | 3.58 |
| BBH(1)-153 | ERS2593851 | 19596872 | 17206221 (88%) | 57.34 | 149.17 | 2.6 | Yes | Cambodian | 97.6 | No | Yes | 3.86 |
| BBH(1)-162 | ERS2593852 | 322082372 | 32342978 (10%) | 72.36 | 102.41 | 1.42 | No | Single copy | 98.5 | No | No | 0.93 |
| HT(1)-144 | ERS2593853 | 16508858 | 8025807 (49%) | 28.52 | 30.73 | 1.08 | No | Single copy | 95.9 | No | No | 1.18 |
| HT(1)-147 | ERS2593863 | 118998744 | 97833898 (82%) | 234.14 | 761.45 | 3.25 | Yes | Cambodian | 98 | No | Yes | 3.71 |
| HT(2)-112 | ERS2593864 | 44338048 | 23619451 (53%) | 73.45 | 83.3 | 1.13 | No | Single copy | 97.6 | No | No | 1.20 |
| JHC(1)-208 | ERS2593865 | 381752146 | 294998614 (77%) | 112.34 | 254.12 | 2.26 | Yes | Cambodian | 98.7 | No | Yes | 2.70 |
| JHC(2)-100 | ERS2593855 | 17622060 | 15958894 (91%) | 56.23 | 166.06 | 2.95 | Yes | Cambodian | 97.3 | No | Yes | 3.24 |
| MKH(1)-72 | ERS2593857 | 17668719 | 14101177 (80%) | 57.11 | 53.41 | 0.94 | No | Single copy | 97.2 | No | No | 1.28 |
| MKH(2)-71 | ERS2593866 | 217327926 | 208538370 (96%) | 715.07 | 1746.88 | 2.44 | Yes | Malagasy | 98.5 | Yes | No | 3.69 |
| SGH(1)-355 | ERS2593858 | 18782232 | 17776334 (95%) | 75.92 | 104.84 | 1.38 | No | Single copy | 97 | No | No | 1.54 |
| SGH(1)-357 | ERS2593859 | 82415713 | 51869956 (63%) | 106.63 | 469.81 | 4.41 | Yes | Malagasy | 98 | Yes | No | 5.14 |
| SGH(1)-331 | ERS849385 | 17931182 | 16408183 (92%) | 90.97 | 79.85 | 0.88 | No | Single copy | 99.7 | No | No | 1.00 |
| SGH(1)-337 | ERS2593867 | 16492008 | 7883420 (48%) | 29.05 | 79.25 | 2.73 | Yes | Cambodian | 95 | No | Yes | 3.44 |
| SGH(1)-358 | ERS2593868 | 13579936 | 7433474 (55%) | 27.75 | 80.08 | 2.89 | Yes | Cambodian | 95.8 | No | Yes | 3.86 |
| SGH(1)-359 | ERS2593869 | 24113148 | 15808023 (66%) | 46.36 | 118.61 | 2.56 | Yes | Cambodian | 97.9 | No | Yes | 3.51 |
| SGH(2)-103 | ERS2593861 | 33312612 | 29984852 (90%) | 123.54 | 272.61 | 2.21 | Yes | Cambodian | 97.8 | No | Yes | 4.40 |
| SGH(2)-108 | ERS2593871 | 6209652 | 4333558 (70%) | 16.45 | 27.06 | 1.64 | Yes | Cambodian | 64.2 | No | Yes | 2.82 |

§ 'Yes' indicates *PvDBP* duplication; 'No' indicates a single copy.
\* 'Yes' indicates an amplification product was observed in the PCR; 'No' indicates no amplification was observed.

### Quantification of *Plasmodium vivax* DNA

The amount of *P. vivax* DNA in each sample was estimated using the SYBR Green quantitative PCR detection method using primers that targeted the 18S rRNA genes of *P. vivax* [10]. Amplification was conducted in a 20 µl reaction mixture containing 1 µL of genomic DNA, 10 µl 2×SYBR Green qPCR Master Mix (Thermo Scientific), and 0.5 uM primer. The reactions were performed in CFX96 Touch^TM^ Real-Time PCR Detection System (BIORAD), with an initial denaturation at 95°C for 3 min, followed by 45 cycles at 94°C for 30 sec, 55°C for 30 sec, and 68°C for 1 min with a final 95°C for 10 sec. This was followed by a melting curve step of temperature ranging from 65°C to 95°C with 0.5°C increments to determine the melting temperature of the amplified product. Each assay included positive controls of *P. vivax* Pakchong (MRA-342G) and Nicaragua (MRA-340G) isolates, in addition to negative controls including uninfected samples and water. A standard curve was produced from a ten-fold dilution series of the control plasmids to determine the amplification efficiency. Melting curve analyses were performed for each amplified sample to confirm specific amplifications of the target sequence. A cut-off threshold of 0.02 fluorescence units that robustly represented the threshold cycle at the log-linear phase of the amplification and above the background noise was set to determine *Ct* value for each assay. The mean *Ct* value was calculated from three independent assays of each sample. Samples yielding *Ct* values higher than 40 (as indicated in the negative controls) were considered negative for *Plasmodium* species. The amount of parasite DNA in a sample was quantified using the follow equation: Parasite DNA (/uL) = [2 ^Ex(40-*Ct*sample)^/10]; where *Ct* for the threshold cycle of the sample, and E for amplification efficiency; assuming 10,000 18S rRNA copies/mL (i.e., 10 copies/uL) in each *Plasmodium* genome [30].

### PCR diagnosis of *PvDBP* duplications

Two sets of primers were used to examine the prevalence and type of *PvDBP* duplications following published protocols [23,24]. The first set of primers includes AF+AR, BF+BR, and BF+AR [23]. Primers BF+AR amplify a 612-bp product that contains the junction between the *PvDBP* copies in isolates with the Malagasy-type duplications. The second set of primers includes AF2+AR2, BF+BR, and BF+AR2 [24]. Primers BF+AR2 amplify a 736-bp product that contains the junction between the *PvDBP* copies in isolates with the Cambodia-type duplication. This primer pair, in theory, should also amplify a 1584-bp product should there be Malagasy-type duplications. Without duplications, BF+AR and BF+AR2 primers are opposite-facing in the samples and thus do not amplify a product [24].

### Estimation of *PvDBP* copy number by qRT-PCR

*PvDBP* copy number was measured with a SYBR Green qPCR detection method using primers (forward: 5’-AGGTGGCTTTTGAGAATGAA-3’; reverse: 5’-GAATCTCCTGGAACCTTCTC-3’) designed between region II to III of *PvDBP* (PVX_110810). *Plasmodium vivax* aldolase, which is known to be a single-copy gene, was used as an internal reference to calculate gene copy number [31]. Two Cambodian isolates, Pv0430 and Pv0431, which were confirmed previously by whole genome sequencing to contain a single and two *PvDBP* copies [24], respectively, were included in each run as positive controls. Water was used as a no DNA control.

Amplification was conducted in a 20µL reaction mixture containing 1µL of genomic DNA (all samples was standardized to ~50 genomes/µL in each reaction based on 18S qPCR assay; Supplementary Table 2), 10µL 2×SYBR Green qPCR Master Mix (Thermo Scientific, USA), and 0.5µM of each primer. Reaction was performed in CFX96 Touch^TM^ Real-Time PCR System (Bio-Rad) with an initial denaturation at 95°C for 3 min, followed by 45 cycles at 94°C for 30 sec, 55°C for 30 sec, and 68°C for 1 min with a final 95°C for 10 sec [22]. This was followed by a melting curve step of temperature ranging from 65°C to 95°C with 0.5°C increments to determine the melting temperature of the amplified product. Each assay included an internal reference *P. vivax* aldolase as well as the negative controls (uninfected samples and water). Melting curve analyses were performed for each amplified sample to confirm specific amplifications of the target sequence. A cut-off threshold of 0.02 fluorescence units that robustly represented the threshold cycle at the log-linear phase of the amplification and above the background noise was set to determine *Ct* value for each assay. The amplification of the *P. vivax aldolase* and *PvDBP* gene fragments was optimized to obtain similar amplification efficiency. The mean *Ct* value and standard error was calculated from three independent assays of each sample (Supplementary Table 3). *PvDBP* copy number (*N*) was calculated with the equation previously used for *Pvmdr*1 copy number estimation [31] as follow: *N* = 2^ΔΔCt±SD^, where ΔΔCt = (Ct_*pvaldo*_-Ct_*pvdbp*_)-(Ct _*pvald*o cal_-Ct_*pvdbp* cal_). The Ct_*pvaldo*_ and Ct_*pvdbp*_ are threshold cycle values for the *P. vivax aldolase* and *PvDBP* genes respectively, whereas Ct_cal_ is an average difference between Ct_*pvaldo*_ and Ct_*pvdbp*_ obtained for the positive control containing a single copy of the *P. vivax aldolase* and *PvDBP* gene fragments (i.e., the Cambodian isolate Pv0430 in this case). The SD value is a standard deviation calculated as follow: 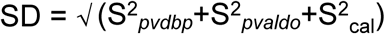 where S_*pvdbp*_ and S_*pvaldo*_ are the standard deviations from the average Ct calculated for three replicates in the *P. vivax aldolase* and *PvDBP* gene amplifications. S_cal_ is an average standard deviation of the ΔCt values for the calibrator. The *PvDBP* copy number of each sample was assessed twice for validation.

### Duffy blood group genotyping

For all *P. vivax* positive samples, a 1,100-bp fragment of the human Duffy blood group antigen gene that encompasses the GATA1 transcription factor-binding site of the gene promoter was amplified using published primers [12]. PCR reaction contained 20μl DreamTaq PCR Mastermix, 1μl DNA template, and 0.5μl each primer. PCR conditions were: 94°C for 2-min, followed by 35 cycles of 94°C for 20s,58°C for 30s, and 68°C for 60s, followed by a 4-min extension. PCR products were sequenced and the chromatograms were visually inspected to determine the *Fy* genotypes (see Duffy blood group nomenclature in [32]).

### Correlation analyses

A one-tailed t-test was used to test for the significance of differences in *PvDBP* copy number between symptomatic and asymptomatic *P. vivax* infections, as well as among Duffy genotypes. In addition, we calculated Pearson's correlation coefficient (*r*^2^) in R for the associations of *PvDBP* copy number with parasitemia and age among the clinical samples. Samples were also stratified by gender and ethnicity to test if there was a significant difference in *PvDBP* copy number. Malaria symptoms including fever, chills, malaise, fatigue, muscle/joint pain, headache, nausea, vomit, diarrhea, abdomen pain, loss of appetite, and respiratory difficulty (dependent variables) were recorded as presence or absence for each of the patients. Principle component analyses were performed to show the variation of all variables among the samples with different *PvDBP* copy number using the logistic PCA function in R. A 3-dimensonal PCA plot was constructed with the plot3d function from the rgl package.

## Results

### *PvDBP* duplications are common in Ethiopian *P. vivax* infections

To establish whether Cambodian or Malagasy-type *PvDBP* duplications were present in the sample population, whole genome sequences were obtained for a subset of 20 Ethiopian *P. vivax* samples using Illumina sequencing (Table 1). For these 20 samples, 13 to 381 million 150 bp pair-end reads were generated and 10-96% of these reads was mapped to the *P. vivax* reference genome PVP01. The average *P. vivax* genome coverage was 16-715× with over 95% of the genome covered by at least 15 reads in the majority of the samples. The average *PvDBP* coverage was 27-1746×, of which the sequence coverage of the *PvDBP* region was clearly higher than the flanking regions in some samples (Supplementary Figure 1). We further confirmed and identified the type of *PvDBP* duplications by mapping the paired-end reads in a tail-to-tail orientation, respectively, on the Malagasy M15 [23] and Cambodian PV0431 [24] reference genomes. Based on whole genome sequences of the duplication break points, two out of the 20 *P. vivax* samples contained the Malagasy-type and 11 contained the Cambodian-type duplications. The remaining had a single copy *PvDBP* (Table 1).

**Figure 1.**
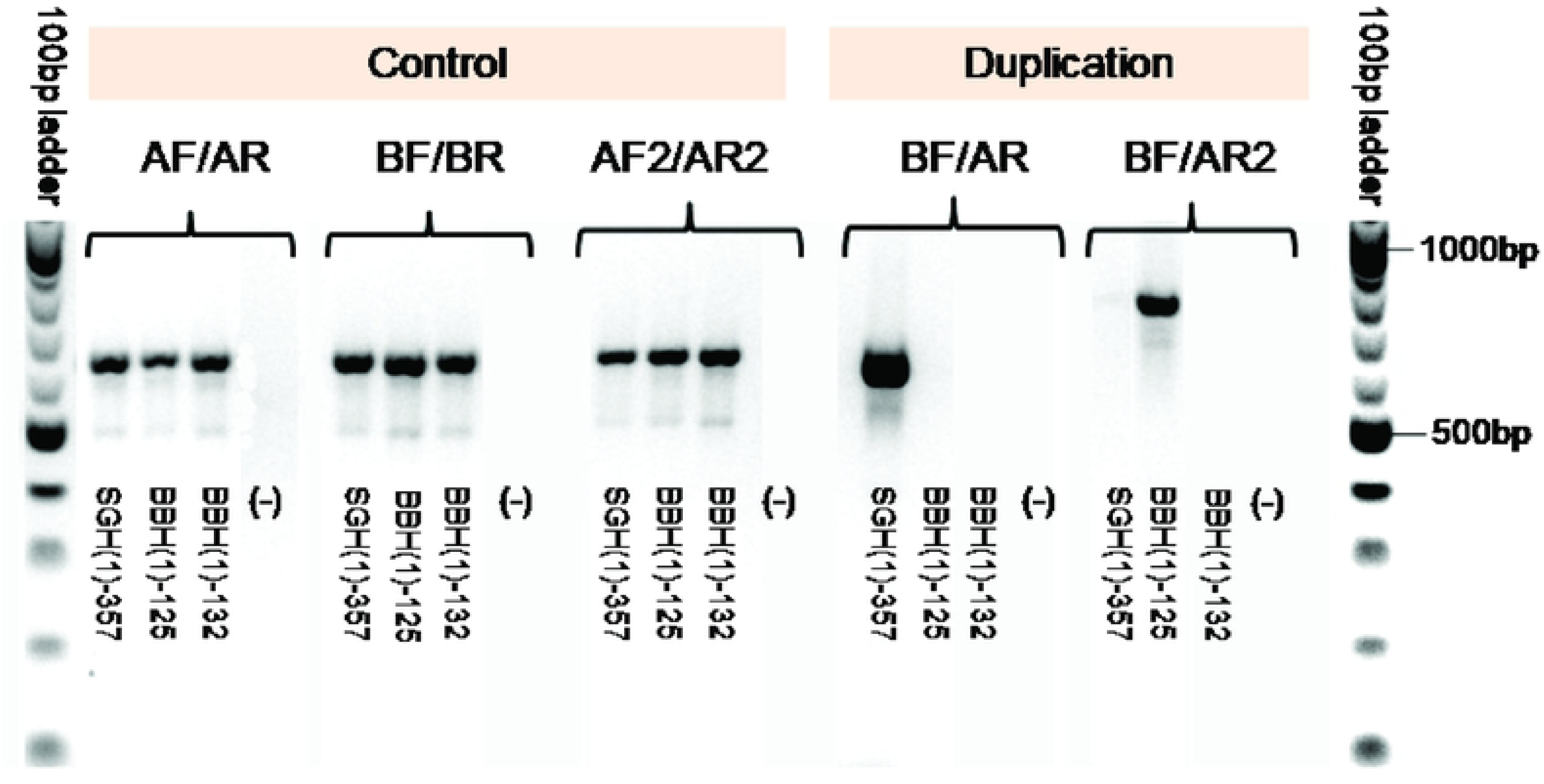
Ethiopian *P. vivax* samples contain both the Malagasy-and Cambodian-type duplications. AF/AR, BF/BR and AF2/AR2 are control primers where all samples indicated amplifications except the negative control sample. Sample SGH(1)-357produced a band of ~650bp with BF/AR primers, indicative of a Malagasy *PvDBP* duplication. Sample BBH(1)-125produced a band of ~750bp with BF/AR2 primers, indicative of a Cambodian *PvDBP* duplication. Sample BBH(1)-132showed no amplification with both BF/AR and BF/AR2 primers, indicative of a single *PvDBP* region without any duplications. No bands were observed for DNA-negative control (-) in all amplifications.

To test more extensively for *PvDBP* duplications, we used recently published diagnostic PCR primers [24]. These primers had previously been used to genotype 20 Ethiopian *P. vivax* symptomatic samples [24]. We genotyped an additional 158 (125 symptomatic and 33 asymptomatic) *P. vivax* samples, bringing the total tested samples to 178 across this and the previous study. All samples produced bands of the expected size of 650-700bp with the control primers AF/AR, BF/BR, and AF2/AR2 (Figure 1), indicating the presence of *PvDBP* in all isolates. Among them, 14 (9.7%) showed a fragment of ~600bp with the Malagasy duplication primers BF/AR (e.g., sample SGH(1)-357 in Figure 1), 81 (55.9%) showed a fragment of ~800bp with the Cambodian duplication primers BF/AR2 (e.g., BBH(1)-125 in Figure 1), and 50 (34.5%) showed no amplifications with either pair of primers (e.g., BBH(1)-132 in Figure 1), suggesting either they only contained a single copy of *PvDBP* or a duplication not detected by either primer pair. For the 14 samples that showed amplification with the Malagasy duplication primers BF/AR, two were amplified with the Cambodian primers BF/AR2 giving a ~1,500bp band.

### A quantitative PCR (qPCR) assay reveals higher order copy number variants in some samples

To test whether additional duplication types not detected by these primer pairs were present, we developed a qPCR assay that compared the quantity of *PvDBP* products to that of a known single copy gene control, *P. vivax* aldolase. To validate the assay, we compared *PvDBP* copy numbers estimated by qPCR with those estimated by PCR diagnostic primers (BF/AR and BF/AR2) and from whole genome sequencing for the 20 samples that had been whole genome sequenced (Table 1). There was a significant correlation between these two metrics (Figure 2A), confirming that the qPCR assay was measuring changes in copy number. For 50 samples that showed no amplification with primers BF/AR and BF/AR2, indicative of a single copy gene, qPCR copy number estimates ranged from 0.85-1.77 (a median value of 1.17±0.26; Figure 2B). Six of these samples had also been whole genome sequenced, and had fold-coverage of *PvDBP* ranging from 0.88-1.42, confirming the presence of only a single *PvDBP* copy (Table 1; Figure 2B). We therefore used qPCR estimates of >1.9 to score samples containing multiple *PvDBP* copies.

**Figure 2.**
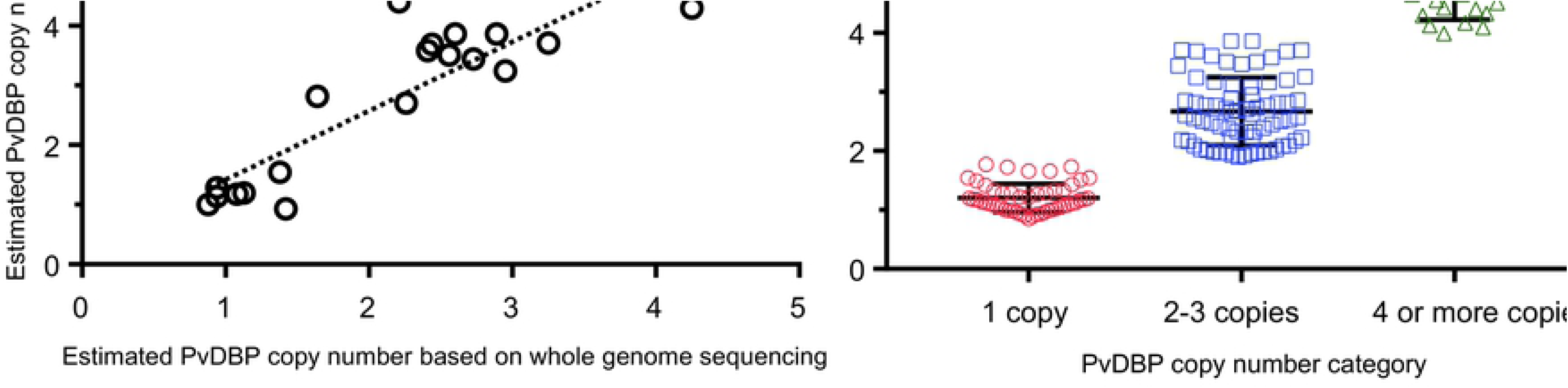
Comparison of *PvDBP* copy number estimated using different approaches. (A) A scatter plot showing *PvDBP* copy number estimated by quantitative real-time polymerase chain reaction (y-axis) and by *PvDBP* coverage in whole genome sequencing (x-axis) of 20 symptomatic *P. vivax* samples. Regression coefficient and *P*-value were indicated. (B) A dot plot showing *PvDBP* copy number detected by quantitative real-time PCR of all 145 symptomatic *P. vivax* samples by three different copy number categories. The geometric median and standard deviation of the data were indicated.

Among the 145 symptomatic samples, there were 95 samples with *PvDBP* qPCR copy number estimates >1.9, and all of these also showed amplifications with BF/AR or BF/AR2 primers, confirming the presence of more than one *PvDBP* copy. qPCR estimates for these samples ranged from 1.90-6.91 (Fig 2B), suggesting that some samples might contain higher order amplifications. We subdivided these samples into two categories: (1) samples with value <3.5 were defined as 2-3 *PvDBP* copies and (2) samples with value ≥3.5 were defined as ≥4 *PvDBP* copies. By this definition, 71 samples had 2-3 *PvDBP* copies (48.9%; median value of 2.42±0.49) and 24 had ≥4 *PvDBP* copies (16.5%; median value of 4.40±0.89; Figure 2B). This is to our knowledge the first report of higher order *PvDBP* copy numbers in a large number of *P. vivax* isolates. *PvDBP* fold coverage estimated from whole genome sequences corroborated the qPCR results in 16 out of the 20 samples. In four samples, qPCR data estimated >4 *PvDBP* copies whereas whole genome sequencing data estimated 2-3 *PvDBP* copies (Supplementary Table 4A). For these four samples, we used the fold-coverage based on the whole genome sequences to score their copy number category. In total, 95 of 145 symptomatic samples tested (65.5%) had more than a single copy of *PvDBP*.

For the 33 asymptomatic samples, two (6.1%) were found with multiple *PvDBP* copies (Fig. 3; Supplementary Table 4B). The qPCR data was further confirmed with the PCR diagnostic primers BF/AR and BF/AR2 for all the 33 asymptomatic samples (Supplementary Table 4B). No amplification was observed in 31 samples, indicative of the absence of duplication. In two asymptomatic samples, amplification was observed with primers BF/AR2, indicative of the Cambodian-type duplication. Two *PvDBP* copies were detected by qPCR in these two asymptomatic samples. A ten-fold higher rate of multi-copy *PvDBP* infections was observed in symptomatic patients compared to the asymptomatic volunteers (*P*<0.001).

**Figure 3.**
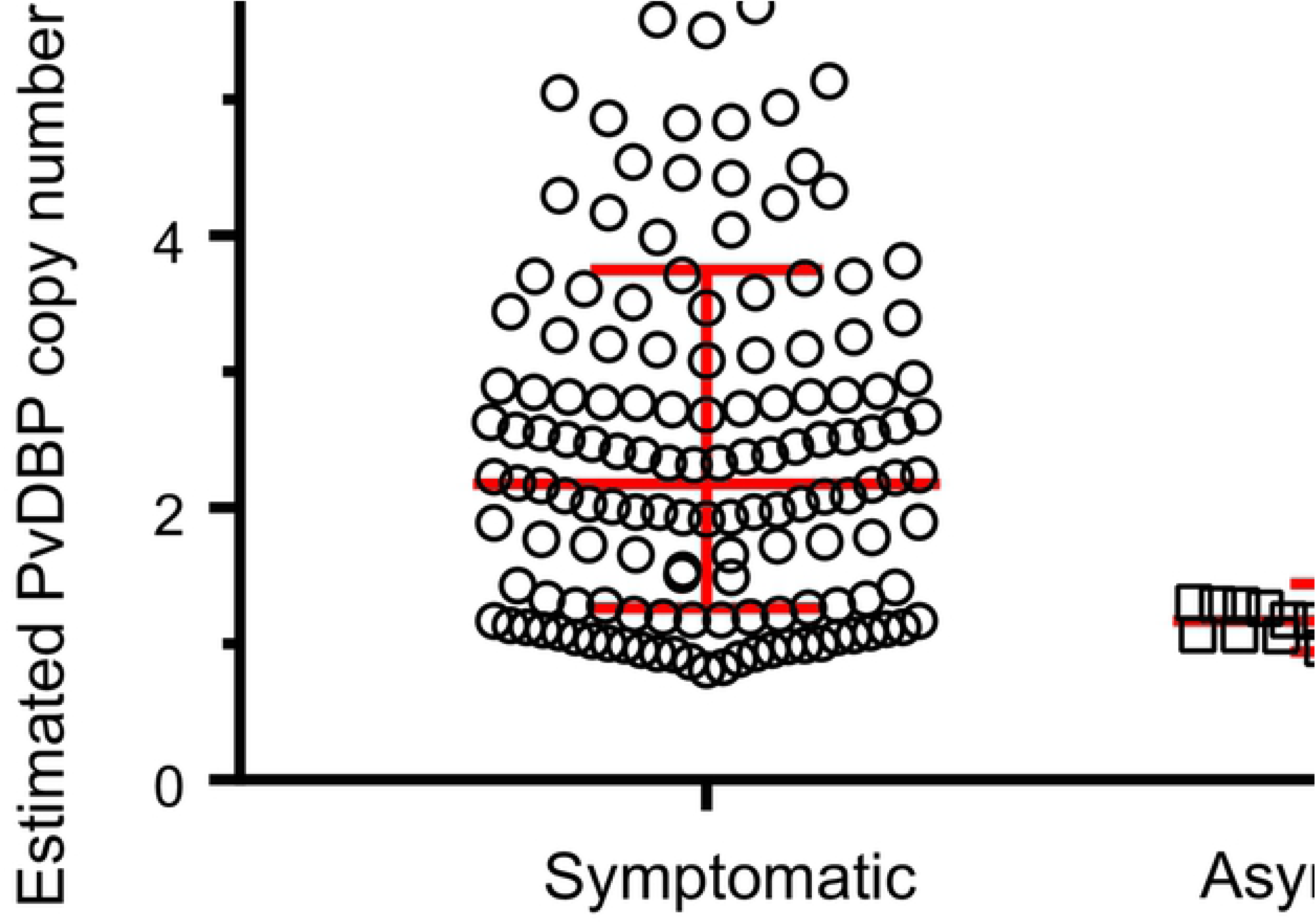
Comparison of *PvDBP* copy number in symptomatic and asymptomatic *P. vivax* samples. *PvDBP* copy number was estimated by qPCR. The geometric median and stand deviation of the data were shown for each type of infection.

### *PvDBP* copy number expansion correlates with malaria symptoms

Based on *DARC* gene sequences, 100 of 145 (69%) symptomatic samples were Duffy-null heterozygote (FyA/B^ES^orFyB/B^ES^), two (1.4%) were Duffy-null homozygote (FyB^ES^/B^ES^), 18(12.4%) were Duffy-positive homozygote (FyA/A orFyB/B) and 25 (17.2%) were Duffy-positive heterozygote (FyA/B or FyB/X^2^; Supplementary Table 5). No significant difference was observed in *PvDBP* copy number among these Duffy genotypes, with the exception of Duffy-null homozygote where the sample size was too small to generate significant comparisons (Figure 4). The two asymptomatic samples detected with two *PvDBP* copies were both Duffy-null heterozygote (FyB/B^ES^). The remaining samples with a single *PvDBP* copy comprised both Duffy-null heterozygote and Duffy-positive homozygote and heterozygote (Supplementary Table 5).

**Figure 4.**
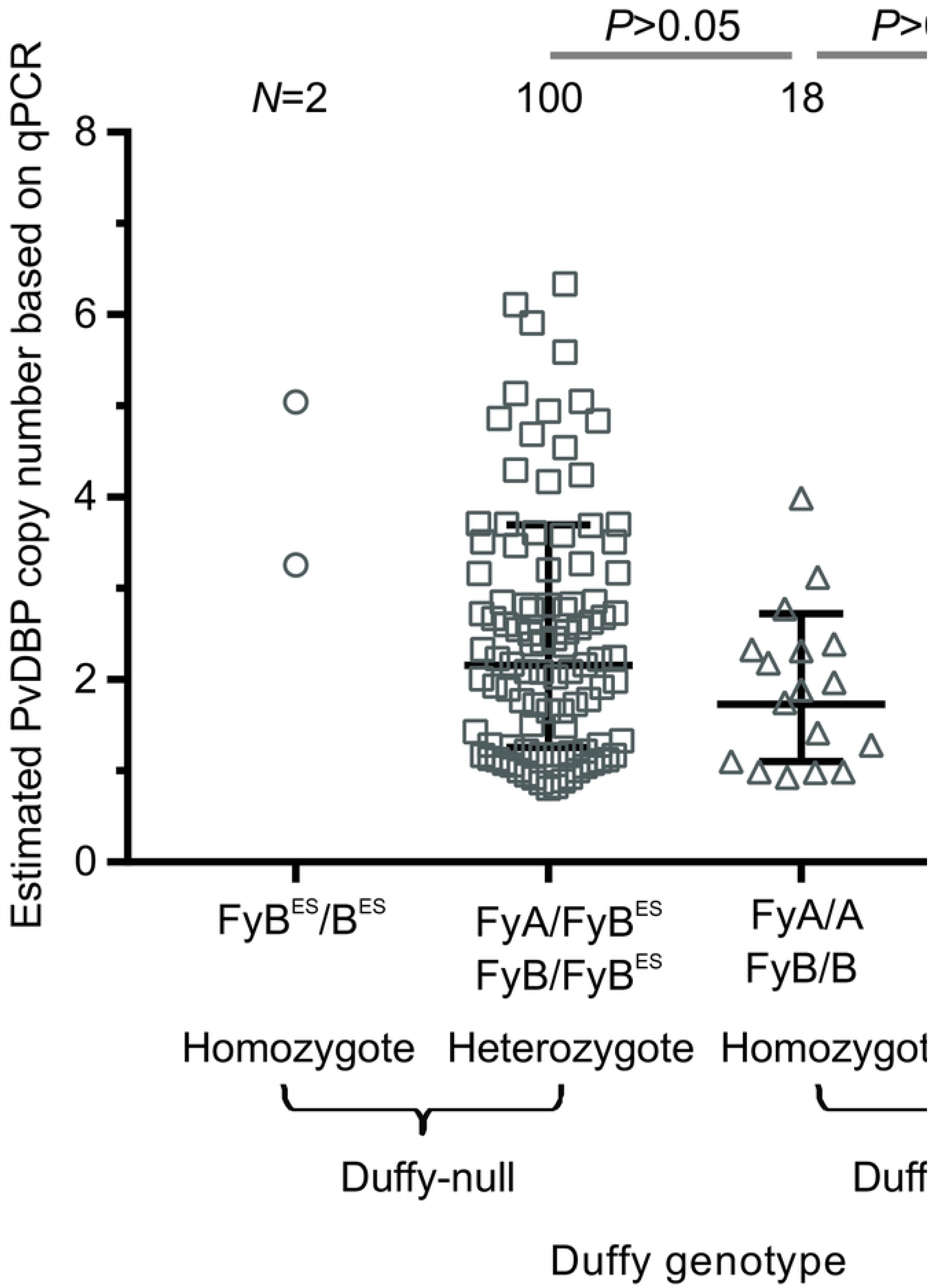
Comparison of *PvDBP* copy number among symptomatic samples with different Duffy genotypes. These included Duffy-null homozygotes (*FyB^ES^*/*B^ES^*), Duffy-null heterozygotes (*FyA*/*B^ES^* or*FyB*/*B^ES^*), Duffy-positive homozygotes (*FyA*/*A* or *FyB*/*B*), and Duffy-positive heterozygotes (*FyA/B* or *FyB/X2*). The geometric median and stand deviation of the data were shown for each of the genotypes.

To investigate whether high *PvDBP* copy number increases invasion efficiency of the parasite, the association of parasite densities with *PvDBP* copy number was examined. No significant association was detected between *PvDBP* copy number and parasite densities in the symptomatic samples (Supplementary Figure 2A). We did not examine this correlation in the asymptomatic samples because of the lack of parasitemia data as well as limited sample size. No significant association was detected between *PvDBP* copy number and age, gender, and ethnicity of the symptomatic samples (Supplementary Figs 2B-D). The PCA plot based on the first three principle components reflected 99.5% of the total variation from 12 recorded malaria symptoms (Figure 5; Supplementary Table 1). No clear cluster was observed among samples with a single or multiple *PvDBP* copies, suggesting that the symptoms of these patients did not relate to *PvDBP* copy number.

**Figure 5.**
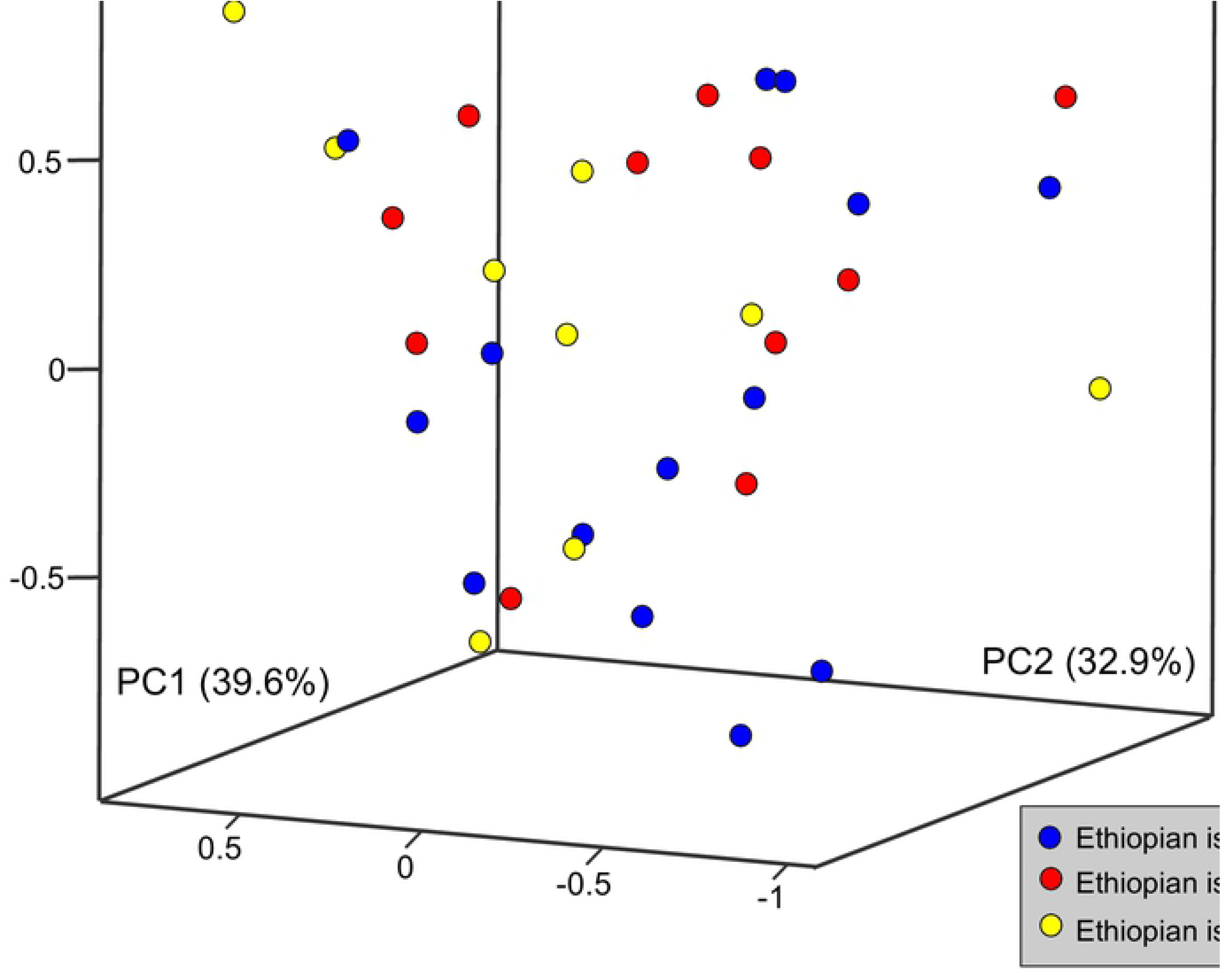
Scatter plot based on principle component analyses showing variation in malaria symptoms among the *P. vivax* patients. *PvDBP* copy number of *P. vivax* observed in each of the patient samples was indicated by different color circles. The first three axes represented 99.5% of the total variation from 12 malaria symptoms. No clear cluster was observed among samples with single or multiple *PvDBP* copies, suggesting that malaria symptoms of these patients did not relate to *PvDBP* copy number. Malaria symptom data are presented in Supplementary Table 3.

## Discussion

*PvDBP* copy number variation has previously been studied primarily using PCR genotyping. In this study, we used a quantitative PCR method for estimating copy number, an approach used in several studies particularly related to human diseases [33]. The qPCR assay outcome was validated against PCR genotyping and whole genome sequencing methods. We are aware that difference in the amount of parasite DNA among samples, particularly that between symptomatic and asymptomatic infections, may influence the quantification of *PvDBP* copy number. Therefore, we standardized the amount of parasite DNA in each reaction based on the results of 18S rRNA quantification prior to *PvDBP* qPCR assay. Also, for each sample we used the single-copy *P. vivax aldolase* gene as internal standard to calibrate and calculate the *PvDBP* gene copy number. Given that the results of *PvDBP* qPCR were consistent with those by PCR genotyping and whole genome sequencing, we are confident that the estimated *PvDBP* copy number was not biased by the amount of parasite DNA in our samples. Quantitative PCR offers a time- and cost-effective approach to analyze a large number of samples, as unlike whole genome sequencing, it can be performed with relatively little DNA, such as dried blood samples routinely taken in malaria studies. It can also detect higher order copy number variants [34], unlike the genotyping PCR methods used in previous studies of *PvDBP* copy number [23,24] that were limited to identifying presence or absence of *PvDBP* duplications.

*PvDBP* duplications detected with specific PCR primers correlated strongly with qPCR estimation of copy number. Samples that showed no amplifications by primers BF/AR (Malagasy-type) [23] or BF/AR2 (Cambodian-type) [24] were all estimated to contain a single copy of *PvDBP* by qPCR. A previous small scale test of only 25 Ethiopian *P. vivax* samples suggested that multi-copy *PvDBP* infections are more common in Ethiopia than in other parts of the world [24]; increasing to 178 samples in this study confirms this finding. Genotyping 178 samples in total revealed that 65% of the Ethiopian isolates contained multiple *PvDBP* copies, which is considerably more than any other *P. vivax* location, as indicated previously in a worldwide study that included only very few samples from Ethiopia [24] - this study provides more statistical robustness to that finding. It is formally possible that qPCR estimates of copy number could be complicated by the presence of mixed infections, where qPCR would generate an estimate that averages all clones present. However, in a previous study, *P. vivax* infections in Ethiopia were found to have a relatively low polyclonality rate (4.3%) based on microsatellite typing [9]. The twenty samples in this study were confirmed as monoclonal by microsatellites prior to whole genome sequencing.

Cambodian-type duplications were five-times more common than the Malagasy-type. Although primers BF/AR2 was expected to produce a 1,584-bp product should there be Malagasy-type duplications [24], only two of the 14 samples that contain the Malagasy-type duplications were detected with such an amplicon, likely due to limited quality of DNA extracted from dried blood on filter papers. The preponderance of Cambodian-type duplications among our samples suggested that this duplication may arise independently in Ethiopia or may be derived from Southeast Asia [35]. It is possible that *P. vivax* with expanded *PvDBP* acquired better fitness and spread through Africa where the majority of populations are Duffy-negative [3]. Though less common, the presence of the Malagasy-type duplications in Ethiopian *P. vivax* may reflect a contemporary gene exchange between populations via human movement.

More than four copies were detected in some of the Ethiopian isolates, higher than that reported in Madagascar (up to two copies) [23] and Southeast Asia (two and three copies) [24,36]. Gene duplication can generate new gene functions or alter gene expression patterns [37]. For examples, in *P. knowlesi* duplication of *PkDBP*-alpha and deletion of *PkDBP*-gamma allow the parasite to invade human erythrocytes that lack surface Neu5Gc, a form of sialic acid *P. knowlesi* requires for binding [38]. In *P. falciparum*, duplications of the *Pfmdr*1 gene resulted in increased resistance to antimalarial drug mefloquine [39–41]. While we did not detect any formal association between *PvDBP* copy number and Duffy negativity, it is worth noting that in other parts of the world such as Cambodia, India, and Brazil where only a small proportion of Duffy-negative individuals live, *PvDBP* expansion was observed with much lower frequency [24]. Although the *PvEBP* gene was shown to be variable in copy number among the Malagasy *P.* vivax [42], we detected only a single copy in the Ethiopian *P. vivax* samples based on whole genome sequences. The functional significance of *PvDBP* expansion merits further investigations through comparison of gene expression patterns and *in-vitro* binding assay of varying *PvDBP* dosage, and study of *P. vivax* isolates from Duffy negative individuals is clearly a high priority.

While we observed no association with Duffy negativity, with the clear caveat that Duffy negative sample numbers were limited, we did observe a statistically significant higher number of *P. vivax* isolates with multiple *PvDBP* copies in symptomatic infections compared to asymptomatic infections. It is therefore possible that increased expression of *PvDBP* as a result of gene expansion may play a role in overcoming the immune response developed by the infected individuals. Further study with higher numbers of asymptomatic samples is needed to draw a definite conclusion. Moreover, symptoms and parasitemia of the samples were measured only at the initial stage of the infection rather than at various follow-up time intervals. Thus, it is yet unclear if high *PvDBP* copies of the parasites will increase severity or duration of *P. vivax* infections. Given that age, gender and ethnicity did not correlate with *PvDBP* copy number, parasites with single or higher *PvDBP* copies could all cause infection equally within the host population.

To conclude, an exceptionally high prevalence of *PvDBP* expansion was observed among Ethiopian *P. vivax* isolates, of which the majority were Cambodian-type duplications, and higher order *PvDBP* copy number variants were identified by both qPCR and whole genome sequencing. Duffy-negative heterozygotes did not show a significantly higher *PvDBP* copy number than the Duffy-positives, but symptomatic infections had a significantly higher copy number than the asymptomatic ones. For the symptomatic samples, *PvDBP* copy number was not significantly associated with parasite density, age, gender and ethnicity. The functional significance of common *PvDBP* expansion and the presence of high copy number variants among the Ethiopian *P. vivax* are unclear. Our ongoing investigations focus on *PvDBP* copy number variation in expanded Duffy-negative homozygote individuals. *PvDBP* copy number in homozygous Duffy-negative infections and its correlation with symptoms are yet to be explored.

## Data Availability

Additional information is provided as supplementary data accompanies this paper. Sequence data of this study are deposited in the European Nucleotide Archive (ENA) and the accession number of each sample is listed in Table 1.

## Acknowledgements

We are greatly indebted to the staffs and technicians from Jimma University for field sample collection, the communities and hospitals for their support and willingness to participate in this research. This research was funded by National Institutes of Health (R01 AI050243, U19 AI129326, D43 TW001505 and R15 AI138002) and the Wellcome Trust (206194/Z/17/Z).

## Author contributions

EL, JBH, JCR, LHM and GY conceived and designed the study. EL, DY and MAH collected the samples. EL, RDP, KG, DK, AF and DAJ collected and analyzed the data. EL, JCR and GY wrote the paper.

## Competing Interests

The authors declare no competing interests.

## Supplementary Data

**Supplementary Table 1.** Presence or absence of malaria signs/symptoms among *P. vivax* patients.

**Supplementary Table 2.** Mean *Ct* value of three independent quantitative real-time PCR assays of *P. vivax* 18S rRNA and the estimated parasite genomes per uL of samples including 145 symptomatic patients and 33 asymptomatic volunteers.

**Supplementary Table 3.** Mean *Ct* value of three independent quantitative real-time PCR assays of *PvDBP* and *Pv aldolase* of 145 symptomatic patients and 33 asymptomatic volunteers. Asterisk indicates samples with multiple *PvDBP* copies.

**Supplementary Table 4.** (A) Results of *PvDBP* duplications based on PCR and copy number estimated by quantitative real-time PCR and whole genome sequencing of *P. vivax* in the 145 symptomatic patient samples; (B) Results of PCR diagnosis and copy number estimated by quantitative real-time PCR of *P. vivax* in the 33 asymptomatic volunteer samples.

**Supplementary Table 5.** Duffy genotypes based on *DARC* sequences of (A) the 145 symptomatic patients and (B) 33 asymptomatic volunteers.

**Supplementary Figure 1.** Coverage view showing mapped reads of sample BBH(1)-125 (blue line) with four-fold higher coverage than sample SGH(1)-331 (red line) with respect to *P. vivax* Sal-1 chromosome 6 region containing *PvDBP* (red box) using the Artemis genome browser.

**Supplementary Figure 2.** Association plots showing the non-significant correlation of *PvDBP* gene copy number with (1) parasitemia level, (B) age, (C) gender, and (D) ethnicity of the *P. vivax* samples.

